# Territorial aggression in male bower-building cichlids *Nyassachromis* cf. *microcephalus* is disrupted by exposure to environmental concentrations of fluoxetine

**DOI:** 10.1101/2022.10.25.513719

**Authors:** Paige Lee, Alan Smith, Hannah King, James Gilbert, Lesley Morrell, Domino Joyce

**Affiliations:** Biological Sciences, School of Natural Sciences, University of Hull, Hardy Building, Cottingham Rd, Hull, HU6 7RX, United Kingdom

**Keywords:** fluoxetine, serotonin, aggression, fish, cichlid, sexual

## Abstract

Selective serotonin reuptake inhibitors (SSRIs) are commonly prescribed drugs for treating human depression, but the role of serotonin—often called the happy chemical—is complex, associated with the regulation of appetite, anxiety, aggression, and more. With such psychoactive pharmaceuticals being increasingly detected in aquatic environments and their effects on non-target species an uncertainty, this study seeks to investigate how inhibiting the serotonin pathway with the SSRI fluoxetine affects territorial aggression, using the cichlid fish *Nyassachromis* cf. *microcephalus* as a model. Males of this sand-dwelling Lake Malawi species build bowers to attract females within a lekking system, where male-male competition is intense. Being aggressive in defending against intruders would serve to maximise mating opportunities and reproductive success for the territory holding male. However, after a one-week exposure to fluoxetine at environmentally relevant concentrations, a decline in aggression was observed in the male cichlids. This implies the serotonergic system plays an important role in modulating aggression and therefore sexual selection in cichlid fishes, and that environmental pollution has the potential to disrupt these behaviours.

## 1. Introduction

As the global human population grows, accompanying anthropogenic pressures on aquatic ecosystems and biodiversity also escalate (Häder et al., 2020; Khan, Hou and Le, 2021; Secretariat of the Convention on Biological Diversity, 2014). Habitat degradation may be reflected by low biological integrity and ecosystem function of water bodies (Kaufmann and Hughes, 2006); nutrient pollution could lead to algal blooms and the depletion of oxygen levels (Misra, Chandra and Raghavendra, 2011); and pharmaceuticals present in wastewater discharge (Wilkinson et al., 2022), such as contraceptive pills containing oestrogen, have been reported to disrupt population dynamics in fish (Schwindt et al., 2014). In addition to direct threats to biodiversity, anthropogenic effects influence critical behaviours that may indirectly impact species survival as well. Juvenile gilt-headed seabreams *Sparus aurata* were observed to swim slower and display fewer bursts of speed when exposed to sunscreen-treated water (Díaz-Gil et al., 2017), which would likely hinder them when escaping from predators or competing for resources. Exposure to another pharmaceutical product, selective serotonin reuptake inhibitors (SSRIs)—a type of antidepressant that has been found in aquatic systems—was discovered to reduce aggression in male Siamese fighting fish *Betta splendens* (Kania, Gralak and Wielgosz, 2012; Kohlert et al., 2012; Lynn et al., 2007). Aggression in this species is essential for parental care and territorial defence (Jaroensutasinee and Jaroensutasinee, 2003).

Pharmaceuticals are increasingly being detected in the environment, entering water bodies through channels such as sewage discharge that may still have chemical contaminants and inadequate disposal of unwanted medication (Monteiro and Boxall, 2010; Ruhoy and Daughton, 2008, Wilkinson et al., 2022). Although environmental concentrations are usually low, their presence is still a concern as these drugs have been made to be effective at low doses (Arnold et al., 2014; Brodin et al., 2014; Metcalfe et al., 2010; Silva et al., 2012) and there is a significant gap in knowledge regarding their uptake by non-target species in aquatic ecosystems as well as the subsequent effects (Boxall et al., 2012; Fent, Weston and Caminada, 2006; Saaristo et al., 2018). Among these pharmaceutical pollutants are antidepressants in the form of selective serotonin reuptake inhibitors (SSRIs) that target serotonin receptors and pathways in the human brain to modulate mood and anxiety. Evolutionary conservatism has maintained similar mechanisms in other vertebrates and there is growing evidence to show how these psychoactive drugs, even at low concentrations, have a role in influencing animal behaviour, endocrinology, physiology and in the long run, survival (Brooks et al., 2003a; Polverino et al., 2021; Sumpter, Donnachie and Johnson, 2014). With increasing mental health issues in the growing world population (Srivastava, 2009) that require treatment using such medication, there is a pressing need to better understand the impact of SSRI pollution on aquatic wildlife.

Serotonin or 5-hydroxytryptamine (5-HT) has wide-ranging and complex effects on fish (Brooks et al., 2003b; Bacqué-Cazenave et al., 2020; Eisenreich, Greene and Szalda-Petree, 2017; Martin et al., 2017; Prasad, Ogawa and Prahar, 2015), including the alteration of aggressive behaviour (Perreault et al., 2003; Lepage et al., 2005; Zubizarreta et al., 2012). SSRIs temporarily increase the amount of serotonin available by blocking the reuptake of serotonin into neurons (Stahl, 1998). Within cichlids, the SSRI fluoxetine has been found to disrupt the endocrine system and reproduction in *Amatitlania nigrofasciata* and *Cichlasoma dimerus* (Dorelle et al., 2017; Latifi, Forsatkar and Nematollahi, 2015), reduce food consumption in *Cichlasoma dimerus* (Dorelle et al., 2020), and reduce the rate of startle and aggressive behaviours in *Astatotilapia burtoni* (Shih, 2017). In Shih (2017), aggression was studied in the capacity of evaluating social status and behaviour. Here, we study cichlid aggression in the context of territoriality, which ultimately has consequences for reproductive success.

Cichlid fishes make up about 10% of all teleost fish species. They are a diverse family of fish well known for being aggressive and territorial (Arnott and Elwood, 2009; Genner, Turner and Hawkins, 1999; Hirschenhauser et al., 2004; Josi and Frommen, 2021). Haplochromine cichlids are popular models in the study of adaptive radiation and speciation (Genner and Turner, 2005; Kocher, 2004; Moser et al., 2018; Seehausen and van Alphen, 1999). The East African Great Lakes are rich with cichlid species that have recently evolved, with no fewer than 200 species inhabiting Lake Tanganyika, and at least 500 species present in each of Lake Victoria and Lake Malawi (Turner et al., 2001). Speciation within these systems occurs at incredibly high rates (Seehausen, 2000; Seehausen, 2006), providing opportunities for the study of the process in its various stages and the mechanisms involved (Kocher, 2004). Selection based on environmental adaptations is a contributing factor to their diversification (Moser et al., 2018; Muschick, Indermaur and Salzburger, 2012; Stauffer Jr. and Gray, 2004), as is selection driven by assortative sexual preferences (Seehausen and van Alphen, 1999; Selz et al., 2016; Stauffer Jr., McKaye and Konings, 2002).

Within any habitat, limited availability of mating and breeding sites demands that males and/or females engage in aggressive behaviours to secure a territory, defend it, and protect their offspring from predators and competing con- and heterospecifics (Danley, 2011; Holder, Barlow and Francis, 1991; Holzberg, 1978). In Lake Victoria, male cichlids were observed to be predominantly aggressive towards males of a similar colour (Seehausen and Schluter, 2004), meaning that male colour polymorphism could be maintained in a population by the selective advantage that comes with a decrease in aggressive interactions. Female preference for males on the basis of male colouration has been confirmed for recently diverged sister species (Selz et al., 2014). Taken together, it is likely that female mate choice and aggressive competition between males are important factors in the diversification of East African haplochromine cichlids (Pauers et al., 2008; Seehausen and Schluter, 2004; Selz et al., 2016). Many haplochromine cichlids breed well under laboratory conditions, offering sustainable populations for *ex situ* research in controlled environments. The present research uses the haplochromine cichlid *Nyassachromis* cf. *microcephalus* as a model to examine the effects of a chemical environmental pollutant on aggression against territory intruders, and review the possible implications for reproduction and speciation.

*N*. cf. *microcephalus* is found in the south of Lake Malawi, and similar to *Nyassachromis microcephalus* which is widely distributed within the lake (Konings, A. & Kazemb, 2018; Trewavas, 1935). While SSRI pollution has not yet been identified to be a problem for Lake Malawi, it provides a model for other affected inland freshwater ecosystems. A zooplanktivorous sand-dweller, *N*. cf. *microcephalus* cluster at selected breeding grounds called leks where males build sand-castles or “bowers” to attract females. They aggressively defend their territory from other males and heterospecific fishes as females visit and select mates. Apart from serving as a site for courtship displays and spawning, these bowers have been suggested to signal the competitive ability and fitness of the defending male (Martin and Genner, 2009). It is costly for males to establish and maintain a territory on large leks as increased competition between males results in less time spent foraging (Young et al., 2009). The benefit for males at larger leks would be increased encounters with females, which tend to exhibit a preference for congregated males (Isvaran and Ponkshe, 2013). In addition to competing with other dominant males, they also have to defend against subordinate males without bowers that may sneak a copulation with females that have chosen the territory-holding male (Magalhaes, Smith and Joyce, 2017). Males therefore have a strong incentive to be aggressive in defending against intruders in their territory as this maximises mating opportunities and reproductive success for the individual. Understanding the role of the serotonin pathway in male behaviour is important for understanding how aggression is modulated in these fish, and could also offer an important insight into the possibility that with the introduction of fluoxetine into their environment, serotonin levels are expected to rise and a decline in aggression is predicted. In this study, we tested whether the response of males towards intruders in their territory was influenced by treatment with the SSRI fluoxetine hydrochloride, both at environmentally relevant and high concentrations.

## 2. Material and methods

### 2.1. Subjects

The study was conducted between May and August 2021 in the aquarium at the University of Hull, using 33 male *Nyassachromis* cf. *microcephalus* descended from wild-caught populations from Lake Malawi in Africa. Sexually mature and dominant, i.e. vibrantly coloured, males were randomly selected from stock tanks to maximise development of territoriality in the experimental tanks. Each individual was anaesthetised in 200mg/L of tricaine methanesulfonate (MS-222) to tank water solution before a small Passive Integrated Transponder (PIT tag) was inserted into its abdominal cavity, and measurements of standard length (nose to caudal peduncle) and weight were taken.

Subjects were then placed in separate experimental tanks of dimensions 59cm(L) x 45cm(B) x 39cm(H) and monitored for 24 hours to ensure recovery. Each tank was neighboured on one side by a tank housing six female *N*. cf. *microcephalus* to stimulate development of territoriality; the opposite side of the tank and the back of the tank were covered with a black curtain to maximise standardisation of visual stimulation, and the front of the tank was left clear to enable observation. A brick was placed in each tank to simulate a bower, together with an air driven sponge filter to maintain water quality (see Figure 1).

**Figure 1.**
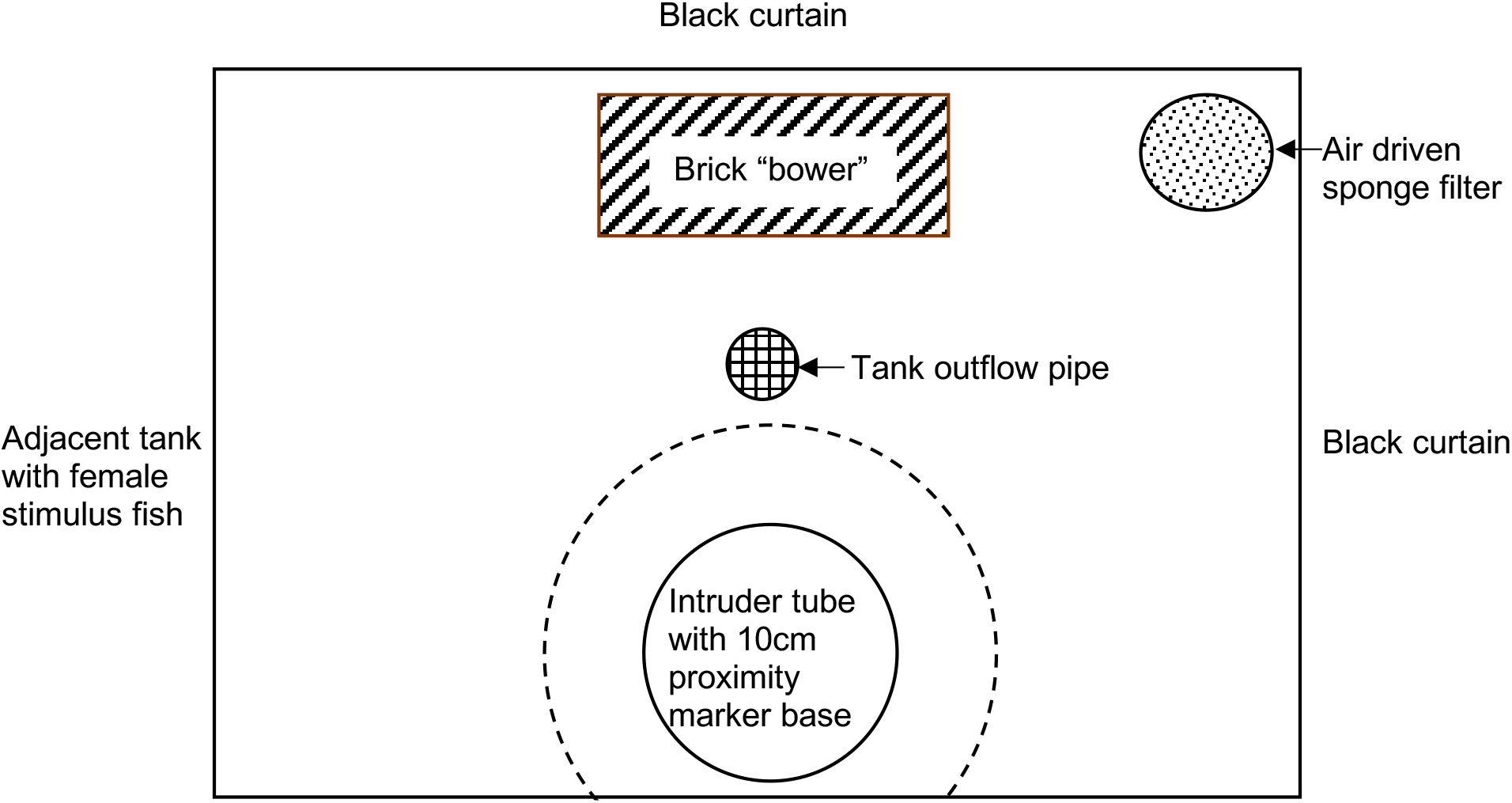
Schematic diagram of experimental tank set up.

During the five days that the subjects were allowed to develop territoriality, the tanks underwent a daily water change of approximately 10% via the aquarium flow-through system. At the end of this period, before fluoxetine treatments were administered, the experimental tanks were taken off the system and the water volume standardised at 70 litres.

The cichlids received a varied diet of ZM (Zebramanagement) flakes, pellets and granular feed once daily. They were fed to satiation except for Day 0 and Day 7 (see following sections) of the experiment. On these days, cichlids were fed 10 pellets and after 10 minutes, remaining pellets were removed and counted. This provided data for assessing appetite change over the treatment period (see Supplementary Materials for results). The aquarium was maintained on a 12:12 light-dark cycle with an ambient temperature of 25°C and water temperature of 24°C.

### 2.2. SSRI treatment

The selective serotonin reuptake inhibitor (SSRI) used in this study was fluoxetine, purchased in hydrochloride form (CAS Number: 56296-78-7) from Fluorochem. Before each treatment period, a fresh stock solution of 540mg/L concentration was prepared by dissolving 54mg of fluoxetine hydrochloride in 100ml of purified water. This stock solution was then further diluted to produce the desired treatment concentrations for each fish.

Each fish was randomly assigned an SSRI treatment using a random number generator (Random.org, 2021a) and dosed on Day 0 by a third person. This ensured that researchers were able to carry out observations on Day 7 without knowing the experimental conditions to avoid bias, i.e. blinded observations. For subjects assigned to the high dose treatment group (n=10), 700μl of the fluoxetine stock solution was added to their tank using a micropipette to produce a concentration of 5.4μg/L. For subjects assigned to the low (environmentally relevant) dose treatment group (n=11), 70μl of stock solution was added to their tank to produce a 0.54μg/L concentration. Subjects assigned to the control group (n=12) had only water added to their tank. These concentrations were selected following reports on fluoxetine presence in the environment (Metcalfe et al., 2010; Silva et al., 2012) and literature on fluoxetine experiments with cichlids (Latifi, Forsatkar and Nematollahi, 2015; Shih, 2017). The high dose group acted as a positive control.

The cichlids were exposed to their respective treatment for seven days, with a refreshment dose administered to the treated tanks every 72 hours (Days 3 and 6). This arrangement was derived from literature reporting the maximum absorption of fluoxetine to occur three days into the exposure period for Japanese medaka (*Oryzias latipes*) (Paterson and Metcalfe, 2008).

The amount of refreshment dose to be administered for each fish was determined following the equation introduced by Barron, Stehly, and Hayton (1990) for calculating the change in amount of chemical in water over time:

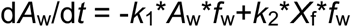

This equation essentially subtracts the amount of drug absorbed by the fish from the amount of drug excreted back into the water. To obtain the former, the uptake rate *k*_1_ was multiplied by the amount of fluoxetine in the water *A*_w_ (i.e. 37.8μg for low dose treatment and 378μg for high dose treatment) and fish weight *f*_w_ (assumed to be constant for each focal fish throughout the experiment). To derive the latter, the elimination rate *k*_2_ was multiplied by the amount of fluoxetine absorbed *X*_f_ (i.e. amount of drug present in fish per gram of body mass) and fish weight *f*_w_.

Barron, Stehly, and Hayton (1990) also provided the formula for calculating *k*_2_ from half-life *t*_½_:

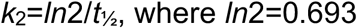

The half-life of fluoxetine in *N*. cf. *microcephalus* was extrapolated from the reported half-life in hybrid striped bass (*Morone saxatilis* x *M. chrysops*) (Gaworecki and Klaine, 2008) and Japanese medaka (Paterson and Metcalfe, 2008), which then allowed us to calculate *k*_2_.

From the comprehensive analysis that Winder et al. (2009) presented of the amount of fluoxetine and its metabolite, norfluoxetine detected in sheepshead minnows (*Cyprinodon variegatus*) after 24, 48 and 72 hours of exposure to the drug, the amount of fluoxetine present in *N*. cf. *microcephalus* after 72 hours *X*_f_ was extrapolated. Together with the *k*_2_ constant derived, the uptake rate of fluoxetine *k*_1_ was calculated following the equation from Barron, Stehly, and Hayton (1990):

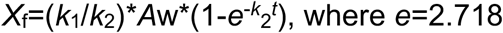

Prior to each refreshment dose, water tests for pH, ammonia and nitrite were conducted to ensure adequate water quality in the absence of automatic water changes whilst being off the aquarium system. Water that had evaporated was also replaced at this point to maintain water volume of 70 litres.

### 2.3. Experimental procedure

At the start of each testing day, two male *N*. cf. *microcephalus* were caught from the stock tanks and placed in a portable container in preparation to serve as intruders for the trials. These individuals were selected based on an estimated colour and size match to the focal fish being trialled on the day (maximum of 10% difference in standard length). Intruder fish were not reused as focal fish.

Before each trial began, a cylindrical tube—made from clear flexible plastic sheeting (14cm diameter tube with 34cm diameter base) and sealed with aquarium-safe silicone—was placed towards the front of the experimental tank (see Figure 1) and filled with untreated water for holding the intruder fish later on. The additional 10cm base radius around the tube (cropped at one section to accommodate tank edge) served as a marker for focal fish proximity to the intruder. The trial commenced once the researcher moved out of visual range of the tank. After 10 minutes, a size-matched intruder was placed into the tube using a net. The trial continued for another 10 minutes after the researcher moved out of view. At the end of each trial, the intruder was first removed from the tube using a net and placed back into the portable container. Water was then siphoned out of the tube, after which, the tube was removed and rinsed thoroughly before being used in another trial. All focal fish were removed from the experimental tanks at the end of the testing day and placed in a designated tank for ‘used’ fish. Intruder fish were also placed in this tank to avoid being accidentally selected as a focal fish at a later date.

Four to ten aggression trials were conducted on each testing day, each lasting for 20 minutes (excluding time taken to introduce intruder fish). Figure 2 shows an experimental trial in progress. The sequence in which the focal fish were trialled was determined using a random sequence generator (Random.org, 2021b). Each trial was recorded using a GoPro HERO6 Black, mounted on the tank using a flexible GoPro tripod and curved extension arm, positioned approximately 20cm in front of the tank. Videos were transferred onto an external hard drive after the experiments and backed up on a cloud storage service (www.box.com) for carrying out behavioural observations.

**Figure 2.**
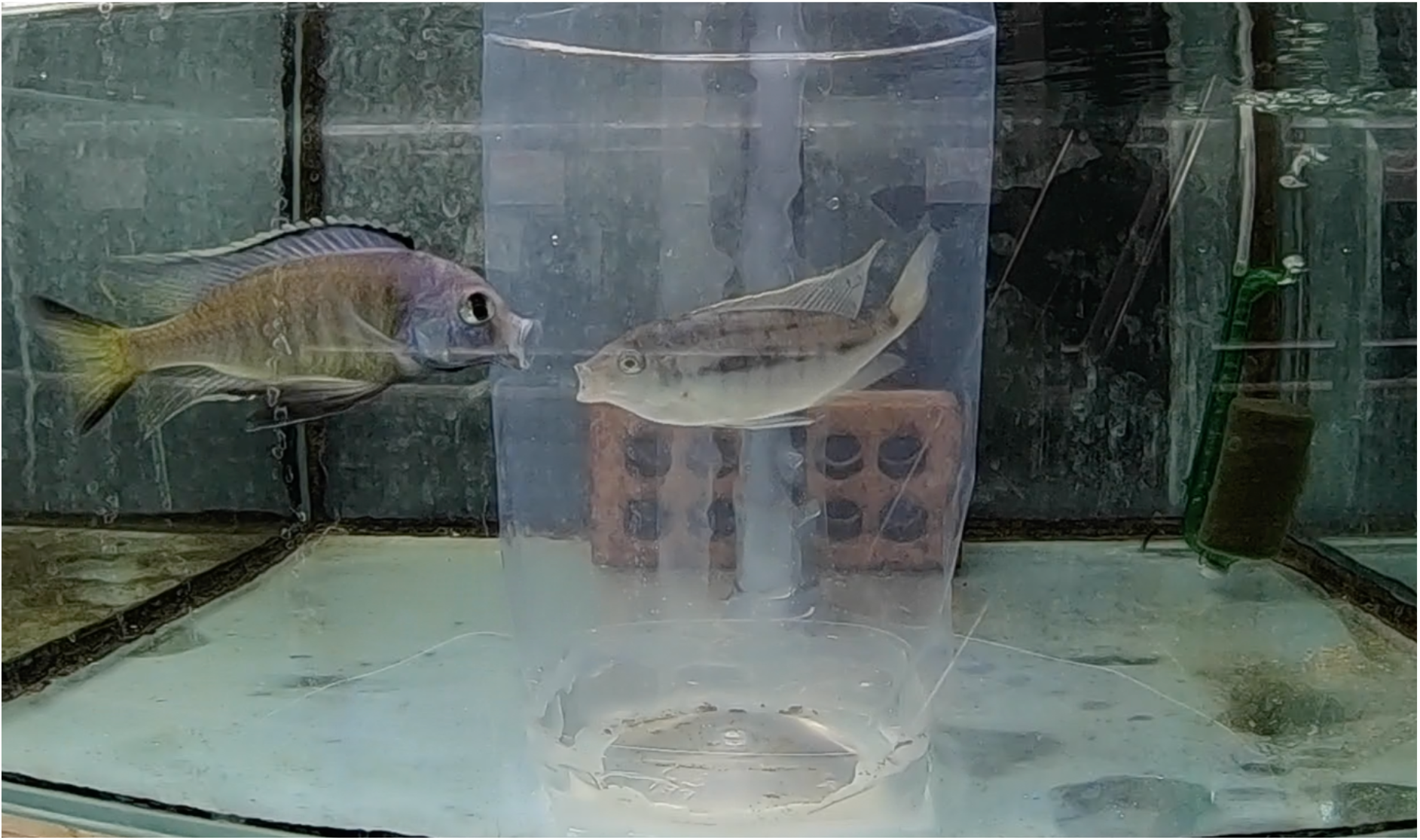
Screenshot of subject 28 displaying “bite” behaviour (see Table 1) during experimental trial.

### 2.4. Behavioural observations

Behaviour was scored using BORIS (Behavioral Observation Research Interactive Software) (Friard and Gamba, 2016), following an ethogram derived from preliminary observations of *N*. cf. *microcephalus* behaviour (Table 1). Subjects were observed for the first 10 minutes of the video, and for another 10 minutes from the 12 minute mark, i.e. a standardised 2 minutes was allowed for intruder introduction. Behaviours recorded during the pre-intruder period served as a baseline of focal fish behaviour with the tube present such that differences in behaviour displayed post-intruder could be attributed to the intruder and not to the novelty of the tube. Data collected was exported as a csv file for conducting statistical analyses, and only at this stage was the experiment unblinded wherein the researchers were informed about the treatment received by each subject.

**Table 1.**
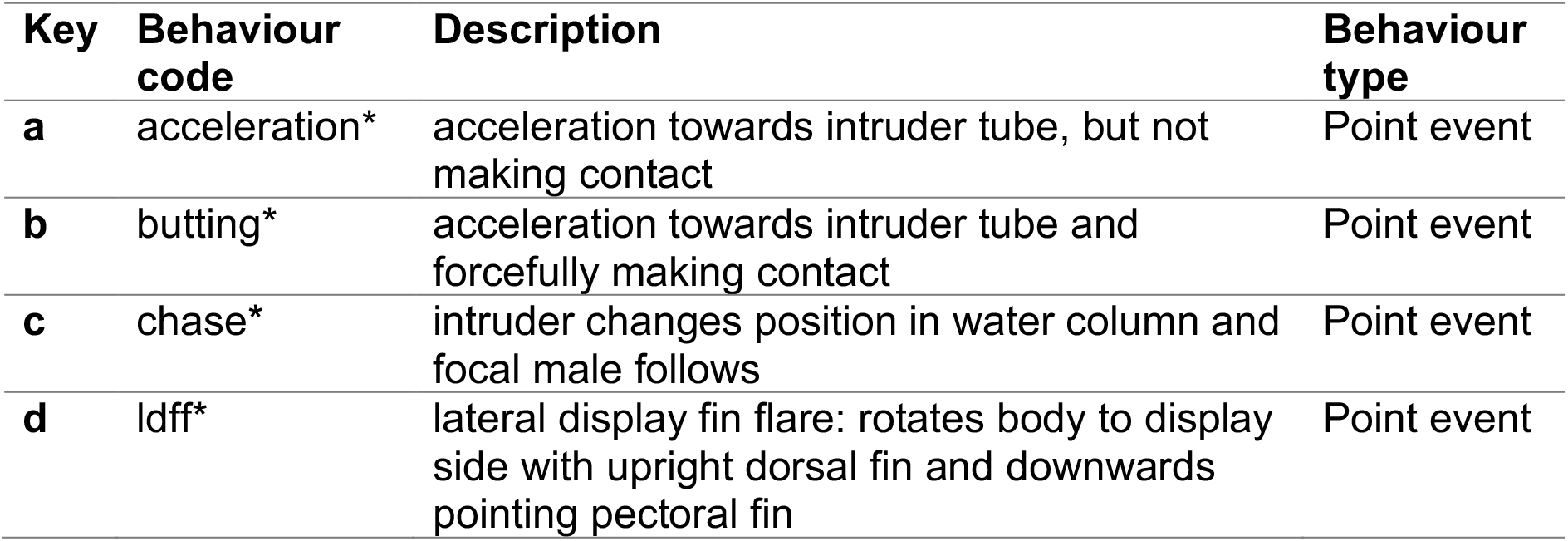
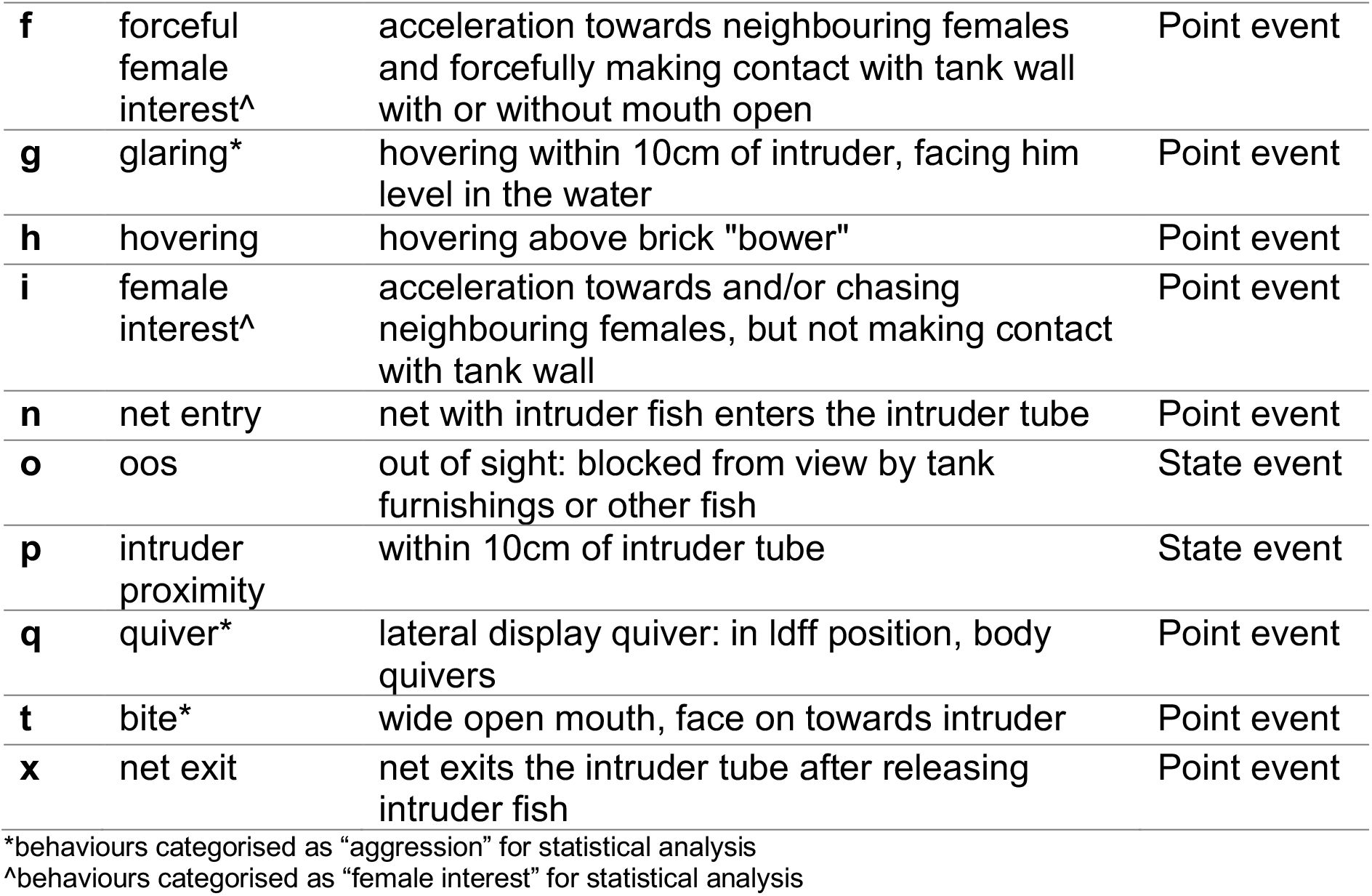
Ethogram used for behavioural analysis of male cichlids.

| Key | Behaviour code | Description | Behaviour type |
| --- | --- | --- | --- |
| a | acceleration* | acceleration towards intruder tube, but not making contact | Point event |
| b | butting* | acceleration towards intruder tube and forcefully making contact | Point event |
| c | chase* | intruder changes position in water column and focal male follows | Point event |
| d | ldff* | lateral display fin flare: rotates body to display side with upright dorsal fin and downwards pointing pectoral fin | Point event |
| **f** | forceful female interest<sup>^</sup> | acceleration towards neighbouring females and forcefully making contact with tank wall with or without mouth open | Point event |
| **g** | glaring<sup>*</sup> | hovering within 10cm of intruder, facing him level in the water | Point event |
| **h** | hovering | hovering above brick "bower" | Point event |
| **i** | female interest<sup>^</sup> | acceleration towards and/or chasing neighbouring females, but not making contact with tank wall | Point event |
| **n** | net entry | net with intruder fish enters the intruder tube | Point event |
| **o** | oos | out of sight: blocked from view by tank furnishings or other fish | State event |
| **p** | intruder proximity | within 10cm of intruder tube | State event |
| **q** | quiver<sup>*</sup> | lateral display quiver: in Idff position, body quivers | Point event |
| **t** | bite<sup>*</sup> | wide open mouth, face on towards intruder | Point event |
| **x** | net exit | net exits the intruder tube after releasing intruder fish | Point event |

<sup>\*</sup>behaviours categorised as "aggression" for statistical analysis
<sup>^</sup>behaviours categorised as "female interest" for statistical analysis

### 2.5. Statistical analyses

Selected behaviours were categorised and their data pooled for analysis (aggression: keys a-d, g, q, t; female interest: keys f,i; see Table 1). Generalised linear mixed models (GLMMs) were fitted for frequency per minute of aggression and duration per minute of intruder proximity to test for treatment as a fixed effect whilst controlling for intruder presence and subject individuality as random effects. A zero-inflated GLMM controlling for subject individuality as a random effect was fitted for frequency per minute of female interest to test the same. A comparison of Akaike Information Criterion (AIC) was used to select the most appropriate models. All statistical analyses were conducted using RStudio (RStudio Team, 2020).

### 2.6. Reproducibility

Videos of all experimental trials are available at: xxx [DOI will be shared when available]; behavioural observations were conducted using these videos on BORIS (Friard and Gamba, 2016) and the project file for this is available at: https://doi.org/10.6084/m9.figshare.21266508. All data generated by and relating to this study, including the raw file produced from scoring behaviour on BORIS are available at: https://doi.org/10.6084/m9.figshare.21231797. All code used in the statistical analyses carried out on RStudio (RStudio Team, 2020) and the resulting output file are available at: https://doi.org/10.6084/m9.figshare.21266493.

### 2.7 Ethical approval

Work was carried out with approval from the University of Hull AWERB and Faculty Ethics Committee, under UK Home Office Project license number P39A1662D.

## 3. Results

Accounting for subject differences and intruder presence as random effects in a Poisson GLMM (model with the lowest AIC value with small sample size adjustment, i.e. delta AICc=1.59; see Table 2), fluoxetine treatment was found to be a significant predictor of aggression (x^2^=802288, df=2, p<0.001).

**Table 2.** Model comparison table for aggression with treatment as a fixed effect, ranked by AICc value.

| Random effects | df | logLik | AICc | delta | weight |
| --- | --- | --- | --- | --- | --- |
| subject + intruder presence | 5 | -124.53 | 260.06 | 0 | 0.533 |
| subject + intruder presence + length | 6 | -124.12 | 261.66 | 1.59 | 0.24 |
| subject + intruder presence + tankid | 6 | -124.53 | 262.49 | 2.43 | 0.158 |
| subject + intruder presence + length + tankid | 7 | -124.12 | 264.16 | 4.1 | 0.069 |
| subject | 4 | -340.11 | 688.87 | 428.81 | 0 |
| intruder presence | 4 | -503.23 | 1015.12 | 755.06 | 0 |

Tukey’s post hoc test using the “glht” function in R indicated a significantly lower frequency of aggressive behaviours in the low dose group compared to the control group (mean difference=0.69 occurrences per minute, p<0.001; see Figure 3), significantly lower frequency of aggressive behaviours in the high dose group compared to control group (mean difference=0.16 occurrences per minute, p<0.001; see Figure 3), and significantly higher frequency of aggressive behaviours in the high dose group compared to the low dose group (mean difference=0.53 occurrences per minute, p<0.001; see Figure 3).

**Figure 3.**
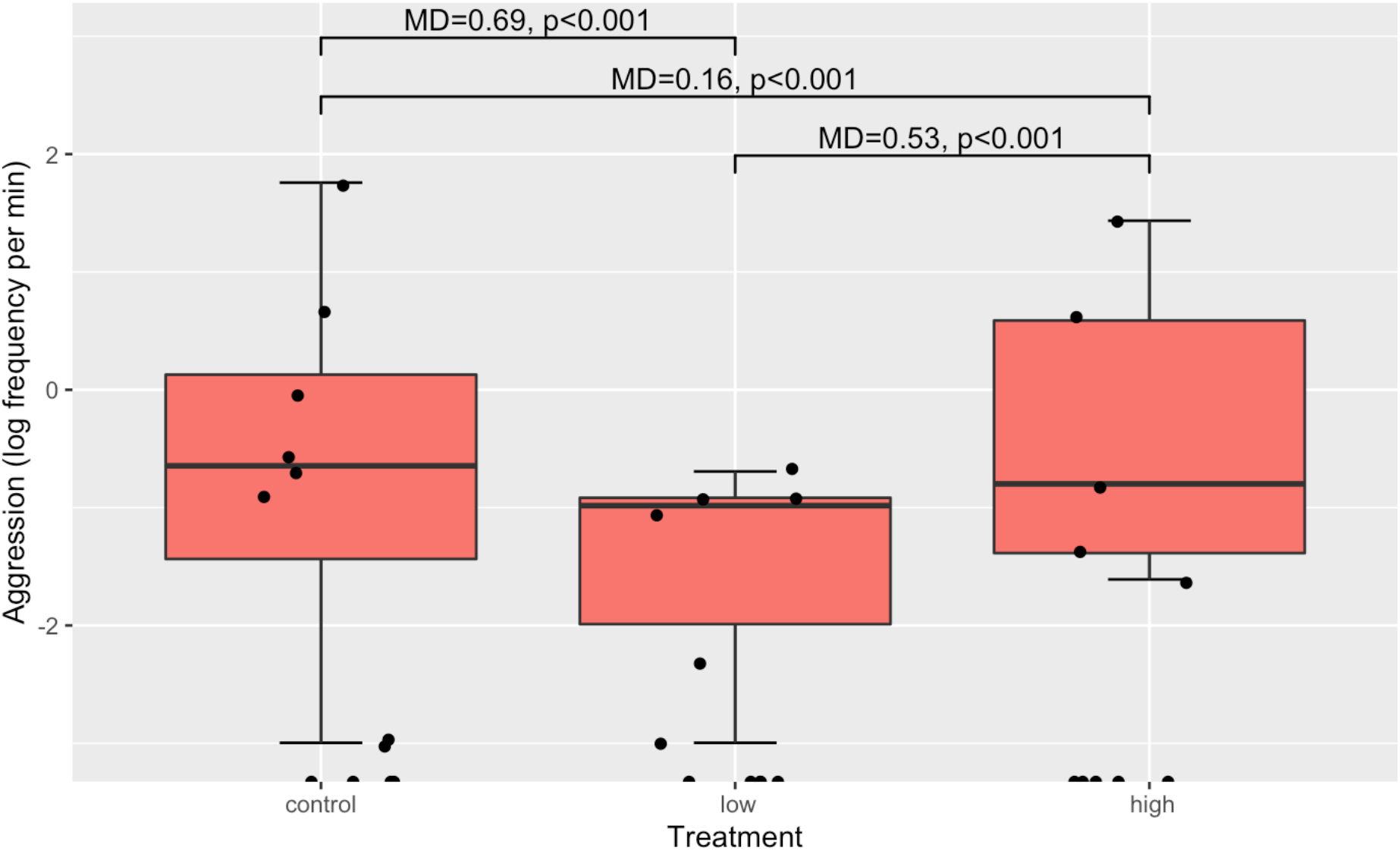
Displays of aggression were significantly fewer in low and high dose groups compared to the control group, and significantly more in the high dose group compared to the low dose group (x^2^=802288, df=2, p<0.001). The round markers represent the frequency of aggressive displays for each subject; see Table 3 for descriptive statistics.

**Table 3.**
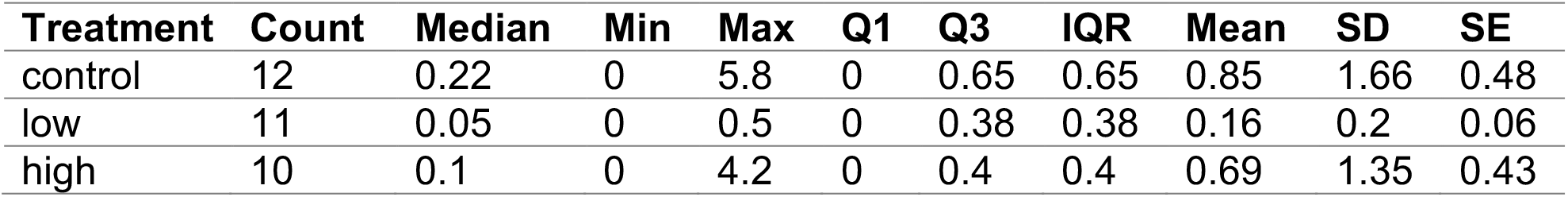
Summary of aggression frequency per minute for each treatment group.

| Treatment | Count | Median | Min | Max | Q1 | Q3 | IQR | Mean | SD | SE |
| --- | --- | --- | --- | --- | --- | --- | --- | --- | --- | --- |
| control | 12 | 0.22 | 0 | 5.8 | 0 | 0.65 | 0.65 | 0.85 | 1.66 | 0.48 |
| low | 11 | 0.05 | 0 | 0.5 | 0 | 0.38 | 0.38 | 0.16 | 0.2 | 0.06 |
| high | 10 | 0.1 | 0 | 4.2 | 0 | 0.4 | 0.4 | 0.69 | 1.35 | 0.43 |

**Table 4.**
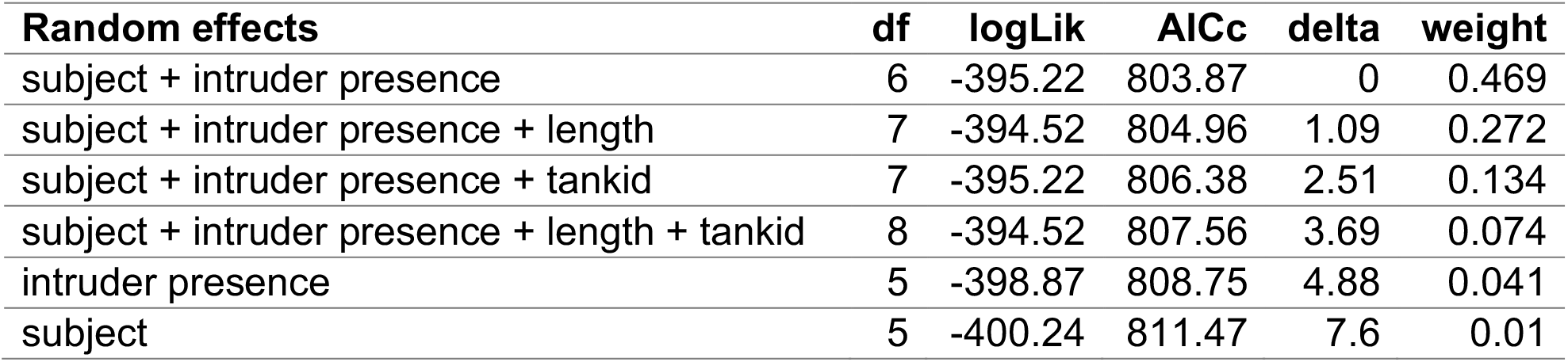
Model comparison table for intruder proximity with treatment as a fixed effect, ranked by AICc value.

| Random effects | df | logLik | AICc | delta | weight |
| --- | --- | --- | --- | --- | --- |
| subject + intruder presence | 6 | -395.22 | 803.87 | 0 | 0.469 |
| subject + intruder presence + length | 7 | -394.52 | 804.96 | 1.09 | 0.272 |
| subject + intruder presence + tankid | 7 | -395.22 | 806.38 | 2.51 | 0.134 |
| subject + intruder presence + length + tankid | 8 | -394.52 | 807.56 | 3.69 | 0.074 |
| intruder presence | 5 | -398.87 | 808.75 | 4.88 | 0.041 |
| subject | 5 | -400.24 | 811.47 | 7.6 | 0.01 |

Accounting for subject differences and intruder presence as random effects in a Gaussian GLMM (delta AICc=1.09; see supplementary Table 4), fluoxetine treatment was not found to be a significant predictor of duration spent in intruder proximity (x^2^=5.51, df=2, p=0.0636; see Figure 4). Fluoxetine treatment was also not found to be a significant predictor of female interest (x^2^=2.09, df=2, p=0.351; see Figure 5) when accounting for subject differences as a random effect in a zero-inflated Poisson GLMM (delta AICc=2.01; see supplementary Table 6).

**Figure 4.**
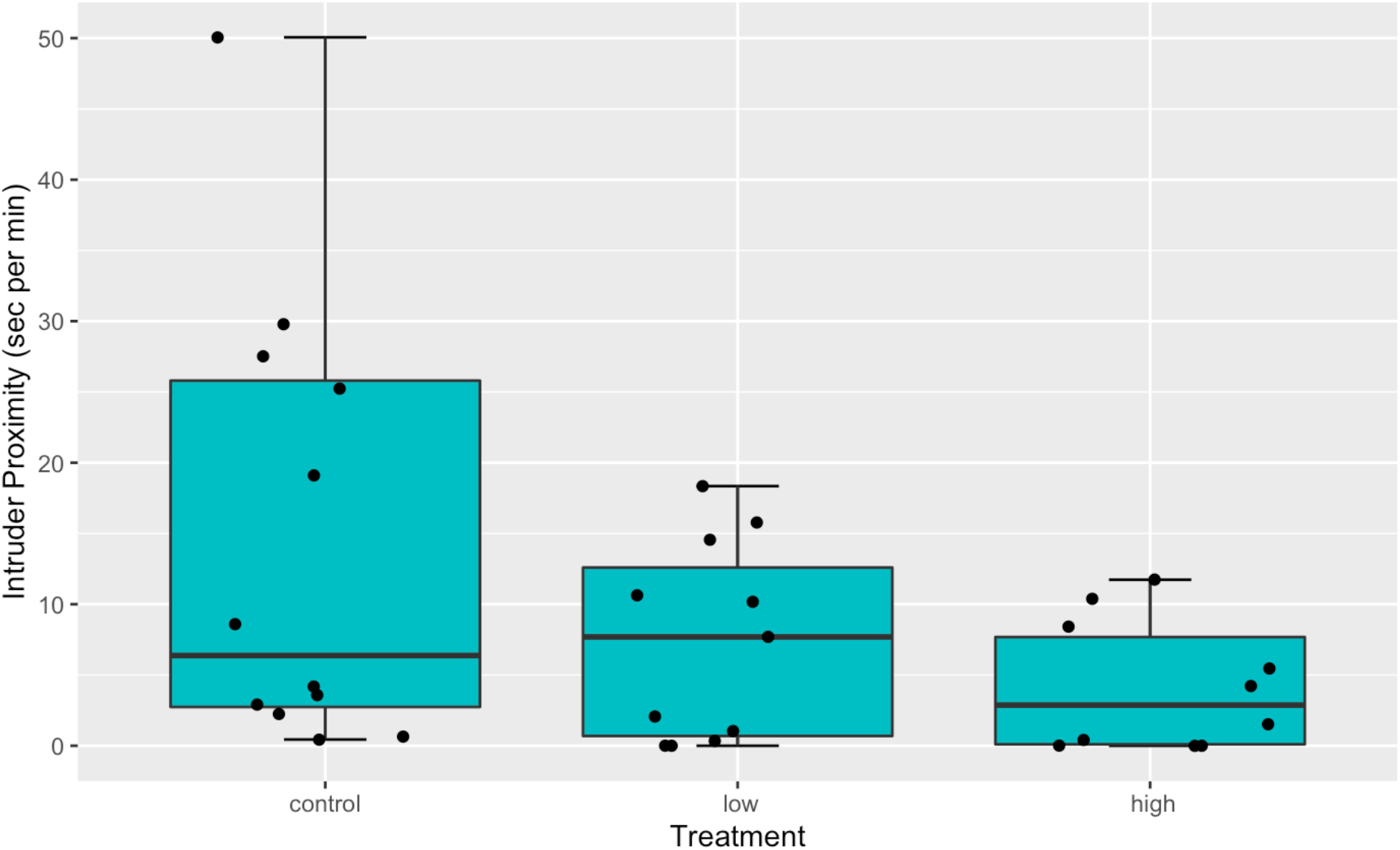
Duration of intruder proximity appears to decrease when treated with higher concentrations of fluoxetine, but did not differ significantly (x2=5.51, df=2, p=0.0636). The round markers represent the duration spent in proximity of the intruder for each subject; see Table 5 for descriptive statistics.

**Table 5.**
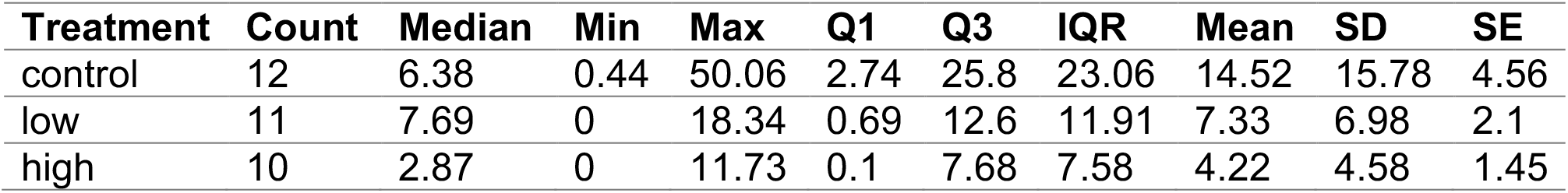
Summary of intruder proximity duration (sec) per minute for each treatment group.

| Treatment | Count | Median | Min | Max | Q1 | Q3 | IQR | Mean | SD | SE |
| --- | --- | --- | --- | --- | --- | --- | --- | --- | --- | --- |
| control | 12 | 6.38 | 0.44 | 50.06 | 2.74 | 25.8 | 23.06 | 14.52 | 15.78 | 4.56 |
| low | 11 | 7.69 | 0 | 18.34 | 0.69 | 12.6 | 11.91 | 7.33 | 6.98 | 2.1 |
| high | 10 | 2.87 | 0 | 11.73 | 0.1 | 7.68 | 7.58 | 4.22 | 4.58 | 1.45 |

**Figure 5.**
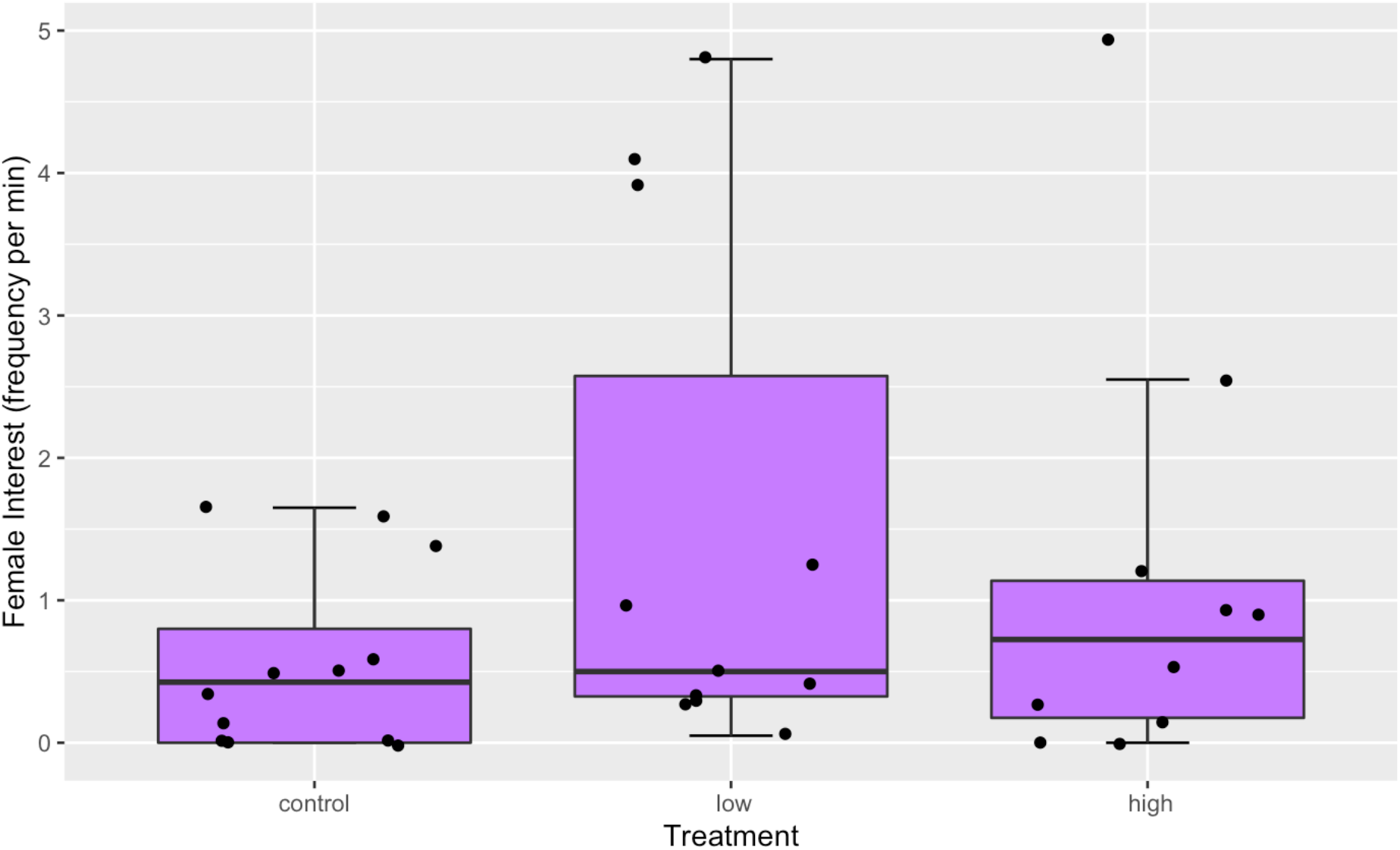
No significant differences and patterns observed in displays of female interest between treatments (x2=2.09, df=2, p=0.351); see Table 7 for descriptive statistics.

**Table 6.** Model comparison table for female interest with treatment as a fixed effect, ranked by AICc value.

| Random effects | df | logLik | AICc | delta | weight |
| --- | --- | --- | --- | --- | --- |
| subject | 4 | -285.70 | 580.1 | 0 | 0.619 |
| subject + intruder presence | 5 | -285.54 | 582.1 | 2.01 | 0.227 |
| subject + intruder presence + length | 6 | -285.54 | 584.5 | 4.44 | 0.067 |
| subject + intruder presence + tankid | 6 | -285.54 | 584.5 | 4.44 | 0.067 |
| subject + intruder presence + length + tankid | 7 | -285.54 | 587.0 | 6.94 | 0.019 |
| intruder presence | 4 | -649.22 | 1307.1 | 727.03 | 0 |

**Table 7.** Summary of female interest frequency per minute for each treatment group.

| Treatment | Count | Median | Min | Max | Q1 | Q3 | IQR | Mean | SD | SE |
| --- | --- | --- | --- | --- | --- | --- | --- | --- | --- | --- |
| control | 12 | 0.43 | 0 | 1.65 | 0 | 0.8 | 0.8 | 0.56 | 0.64 | 0.18 |
| low | 11 | 0.5 | 0.05 | 4.8 | 0.32 | 2.58 | 2.25 | 1.53 | 1.8 | 0.54 |
| high | 10 | 0.72 | 0 | 4.95 | 0.17 | 1.14 | 0.96 | 1.15 | 1.54 | 0.49 |

## 4. Discussion

Studies showing how SSRIs alter aggressive behaviour in fish have slowly been accumulating over the last two decades (Perreault, Semsar and Godwin, 2003; Lepage et al., 2005; Dzieweczynski and Hebert, 2012; Kellner et al., 2018). Findings mostly demonstrate a reduction of aggression with SSRI exposure, and results from the present research support this association. As predicted, male *Nyassachromis* cf. *microcephalus* that had been treated with fluoxetine exhibited less aggression than those that had not. SSRIs have already been established to cause disruptions in fish reproductive fitness, for example decreasing testosterone levels in the male convict cichlid *Amatitlania nigrofasciata* (Dorelle et al., 2017; Latifi, Forsatkar and Nematollahi, 2015; Prasad, Ogawa and Prahar, 2015). In Lake Malawi cichlids, where territoriality is a key component in male mating success and reproduction, the serotonin pathway is likely to play an important role in regulating aggression.

This study found that cichlids exposed to the environmentally relevant concentration of 0.54μg/L performed the least aggressive displays while cichlids that were treated with a higher concentration of 5.4μg/L—and presumably had higher levels of serotonin in their system—were more aggressive (but still significantly less aggressive than cichlids in the control group). This is a similar finding to Kania, Gralak and Wielgosz (2012), in which a group of *Betta splendens* administered with the highest dose of fluoxetine (100μg/g of body weight) were more aggressive than those treated with fluoxetine at 40μg/g of body weight (the least aggressive in their experiment). The mechanisms for this are not clear, but there have been studies on how fluoxetine affects boldness and anxiety related behaviours in recent years that could offer new insights. Mosquitofish *Gambusia holbrooki* expressed anti-anxiety behaviour after being exposed to fluoxetine at high concentrations of 25 and 50μg/L, i.e. an anxiolytic effect (Meijide et al., 2018). Also in 2018, Nielson et al. demonstrated that the SSRI escitalopram increased boldness in zebrafish *Danio rerio* at concentrations as low as 1.5μg/L. Taken together, this could indicate that whilst the primary effect of SSRI is the reduction of aggression, the secondary effects of increasing boldness and reducing anxiety at higher concentrations in turn diminishes inhibitions or influences decision-making, facilitating the expression of agonistic behaviours such that the end-result is reversed. In other words, the aggression displayed in the absence of fluoxetine and aggression displayed at high concentrations of fluoxetine could be driven by different factors. Targeted investigations to address whether similar mechanisms for courtship behaviour would illuminate the complexity of the effects of serotonin and SSRI pollution on animal reproductive behaviour.

This study confirms that fluoxetine exposure decreases territorial aggression in male *N*. cf. *microcephalus* males even at low environmental concentrations. SSRI pollution could therefore cause significant behavioural changes in aggressive species. With a decline in aggression, dynamics in male-male competition and subsequent outcomes would be affected, which could reduce sexual selection. Long term population and species responses to reduced sexual selection could increase extinction risk, because of a dilution of Rowe and Houle’s (1996) “genic capture” process (Martínez-Ruiz and Knell, 2016; Parrett et al., 2019). In this process, condition dependent male traits that derive from additive genetic variance at many loci, and that are subject to female choice, lead to a positive correlation between the sexually selected trait and male condition. Reduced sexual selection therefore reduces selection at these loci. This is likely to be especially acute for species with high reproductive skew such as lekking, bower-building cichlids (Genner et al., 2008) in which dyadic male aggression is known to contribute to phenotypic diversity (Dijkstra et al., 2009). Conversely, it may be that in a habitat where sympatric incipient species of haplochromine cichlids co-exist, there is the possibility that reduced aggression could produce opportunities for novel phenotypes to be maintained in the population until such a time that a female preference could emerge (Magalhaes, Smith and Joyce, 2017; Seehausen and van Alphen, 1999), which may eventually lead to further diversification through runaway sexual selection. Furthermore, cichlid bowers are thought to be honest indicators of quality (Schaedelin and Taborsky, 2006; Martin and Genner, 2009; Taylor et al., 1998) and the serotonergic system is frequently implicated in other relevant behavioural patterns, for example digging behaviour in mice (Deacon, 2006; Hu and Hoekstra, 2017). The role of the serotonergic system in bower building cichlid sexual selection is therefore an exciting avenue of research.

Animal fitness and biodiversity in aquatic ecosystems are already vulnerable to anthropogenic pressures, especially climate change (Baroiller et al., 1995; Bradshaw and Holzapfel, 2008; Walker II et al., 2019). The addition of SSRI pollution as a stressor not only compounds the problem, but the interaction between SSRI effects on behaviour and changes to temperature and photoperiod is also an uncertainty. Temperature has been known to significantly affect chemical toxicity (Holmstrup et al., 2010; Noyes et al., 2009; Roggatz et al., 2019). Whilst environmental concentrations of SSRIs are not considered toxic, it is possible that the efficacy may be influenced by temperature. With warmer waters reported to increase aggression (Kua et al., 2020; Ratnasabapathi and Souchek, 1992; Zubizarreta et al., 2012), examining the interaction between SSRIs and temperature variations could be important. Different climate change scenarios could have significant consequences for this interaction in the context of aggression, and ultimately reproductive behaviour.

## Acknowledgements

We would like to thank Sonia Jennings for technical support in the aquarium and the EvoHull Lab members for discussions throughout the study. We are grateful to Magnus Johnson for statistical advice.

## Supplementary Materials

A Kruskal–Wallis test revealed that appetite change did not differ significantly between fluoxetine treatment groups (x^2^=0.219, df= 2, p=0.896; see Figure 6).

**Figure 6.**
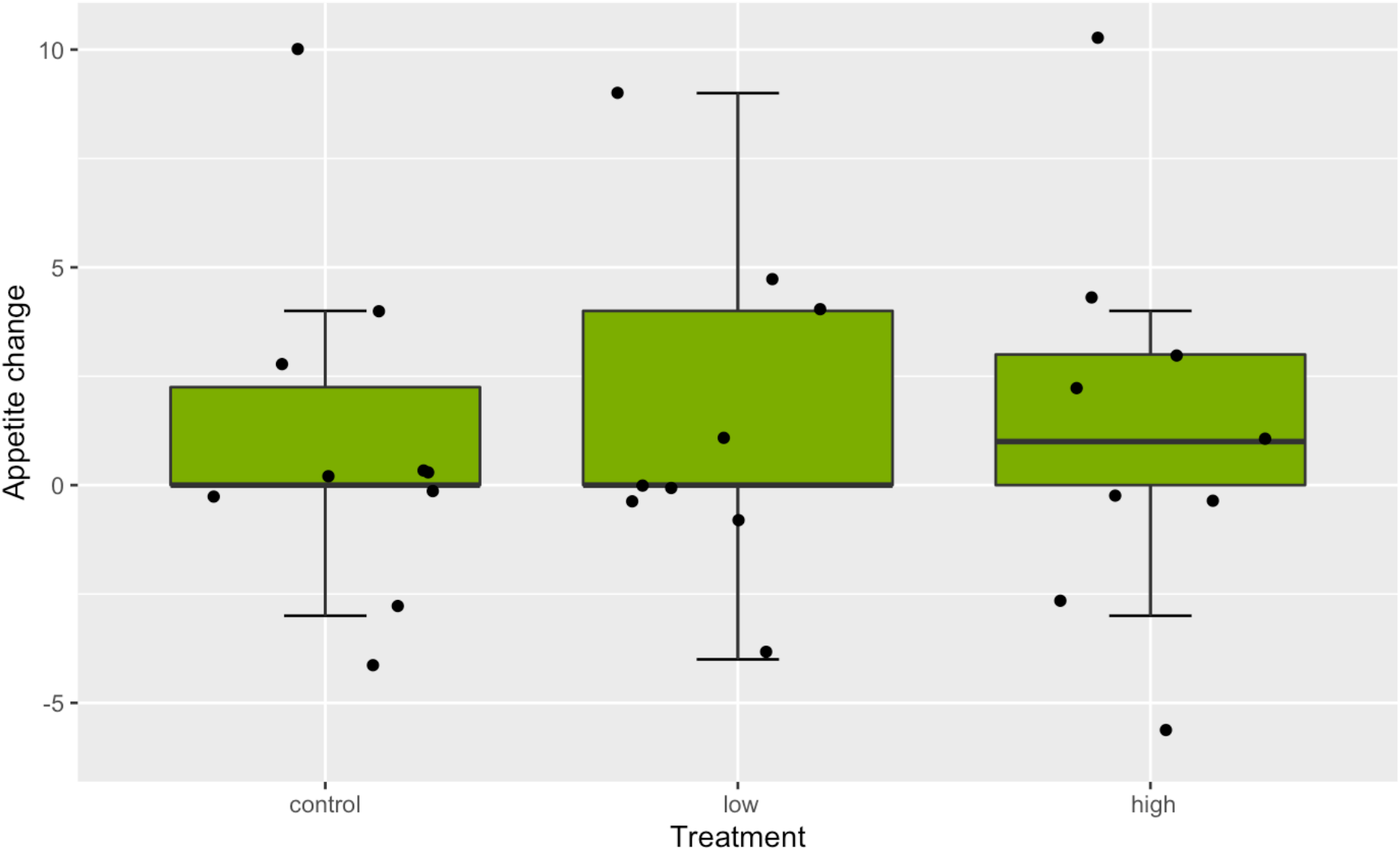
No significant differences observed in appetite change between treatment groups (Kruskal-Wallis x2=0.22, df=2, p=0.896). The round markers represent the change in appetite between experimental Days 0 and 7 for each subject, where negative values indicate an increase in pellets eaten; see Table 8 for descriptive statistics.

**Table 8.** Summary of appetite change for each treatment group.

| Treatment | Count | Median | Min | Max | Q1 | Q3 | IQR | Mean | SD | SE |
| --- | --- | --- | --- | --- | --- | --- | --- | --- | --- | --- |
| control | 10 | 0 | -4 | 10 | 0 | 2.25 | 2.25 | 1 | 3.94 | 1.25 |
| low | 9 | 0 | -4 | 9 | 0 | 4 | 4 | 1.56 | 3.84 | 1.28 |
| high | 9 | 1 | -6 | 10 | 0 | 3 | 3 | 1.22 | 4.49 | 1.5 |

This study did not reveal significant differences in appetite between treatments, contrasting with results from Dorelle et al. (2020) that demonstrated a reduction in appetite and food consumption in *Cichlasoma dimerus* after being injected with 2 or 20μg of fluoxetine per gram of body weight. Extrapolating from Winder et al. (2009), it is estimated that there would have been 0.4 to 0.6μg of fluoxetine per gram of body weight present in the cichlids of this study according to the concentrations they were exposed to; perhaps changes to appetite change may only develop at higher concentrations.

These findings might also be a reflection of how bower-building cichlids prioritise bower defence over eating in the wild (McKaye, 1983), hence appetite was observed to be low across all treatments. Alternatively, data may need to be collected daily to assess feeding trends rather than using only the difference in food consumed on the first and last day of treatment, as was the method used here.

## References

Arnold, K. E., Brown, A. R., Anklew, G. T. and Sumpter, J. P. (2014) Medicating the environment: assessing risks of pharmaceuticals to wildlife and ecosystems. Philosophical Transactions of The Royal Society B, 369(20130569), pp. 1–11. doi:10.1098/rstb.2013.0569.

Arnott, G. and Elwood, R. W. (2009) Gender differences in aggressive behaviour in convict cichlids. Animal Behaviour, 78, pp. 1221–1227. doi:10.1016/j.anbehav.2009.08.005

Bacqué-Cazenave, J., Bharatiya, R., Barrière, G., Delbecque, J-P., Bouguiyoud, N., Di Giovanni, G., Cattaert, D and De Deurwaerdère, P. (2020) Serotonin in Animal Cognition and Behavior. International Journal of Molecular Sciences, 21(1649), pp. 1–23. doi:10.3390/ijms21051649.

Baroiller, J. F., Chourrout, D., Fostier, A. and Jalabert, B. (1995) Temperature and Sex Chromosomes Govern Sex Ratios of the Mouthbrooding Cichlid Fish Oreochromis niloticus. The Journal of Experimental Zoology, 273, pp. 216–223.

Barron, M. G., Stehly, G. R., and Hayton, W. L. (1990) Pharmacokinetic modeling in aquatic animals: I. Models and concepts. Aquatic Toxicology, 17, pp. 187–212.

Boxall, A. B. A., Rudd, M. A., Brooks, B. W., Caldwell, D. J., Choi, K., Hickmann, S., Innes, E., OStapyk, K., Staceley, J. P., Verslycke, T., Ankley, G. T., Beazley, K. F., Belanger, S. E., Berninger, J. P., Carriquiriborde, P., Coors, A., DeLeo, P. C., Dyer, S. D., Ericson, J. F., Gagné, F., Giesy, J. P., Gouin, T., Hallstrom, L., Karlsson, M. V., Larsson, D. G. J., Lazorchak, J. M., Mastrocco, F., McLaughlin, A., McMaster, M. E., Meyerhoff, R. D., Moore, R., Parrott, J. L., Snape, J. R., Murray-Smith, R., Servos, M. R., Sibley, P. K., Straub, J. O., Szabo, N. D., Topp, E., Tetreault, G. R., Trudeau, V. L. and Kraak, G. V. D. (2012) Pharmaceuticals and Personal Care Products in the Environment: What Are the Big Questions? Environmental Health Perspectives, 120(9), pp. 1221–1229. doi:10.1289/ehp.1104477.

Bradshaw, W. E. and Holzapfel, C. M. (2008) Genetic response to rapid climate change: it’s seasonal timing that matters. Molecular Ecology, 17, pp. 157–166. doi:10.1111/j.1365-294X.2007.03509.x.

Brodin, T., Piovano, S., Fick, J., Klaminder, J., Heynen, M. and Jonsson, M. (2014) Ecological effects of pharmaceuticals in aquatic systems—impacts through behavioural alterations. Philosophical Transactions of The Royal Society B, 369(20130580), pp. 1–10. doi:10.1098/rstb.2013.0580.

Brooks, B. W., Foran, C. M., Richards, S. M., Weston, J., Turner, P. K., Stanley, J. K., Solomon, K. R., Slattery, M. and La Point, T. W. (2003a) Aquatic ecotoxicology of fluoxetine, Toxicology Letters, 142, pp. 169–183. doi:10.1016/S0378-4274(03)00066-3.

Brooks, B. W., Turner, P. K., Stanley, J. K., Weston, J. J., Glidewell, E. A., Foran, C. M., Slattery, M., Point, T. W. L. and Huggett, D. B. (2003b) Waterborne and sediment toxicity of fluoxetine to select organisms. Chemosphere, 52, pp. 135–142. doi:10.1016/S0045-6535(03)00103-6.

Danley, P. D. (2011) Aggression in closely related Malawi cichlids varies inversely with habitat complexity. Environmental Biology of Fishes, 92, pp. 275–284. doi:10.1007/s10641-011-9838-7

Deacon, R. (2006) Digging and marble burying in mice: simple methods for in vivo identification of biological impacts. Nature Protocols, 1, pp. 122–124. doi:10.1038/nprot.2006.20.

Díaz-Gil, C., Cotgrove, L., Smee, S. L., Simon-Otegui, D., Hinz, H., Grau, A., Palmer, M. and Catalan, I. A. (2017) Anthropogenic chemical cues can alter the swimming behaviour of juvenile stages of a temperate fish. Marine Environmental Research, 125, pp. 34–41. doi:10.1016/j.marenvres.2016.11.009.

Dijkstra, P. D., Hemelrijk, C., Seehausen, O. and Groothuis, T. G. G. (2009) Color polymorphism and intrasexual competition in assemblages of cichlid fish. Behavioral Ecology, 20(1), pp. 138–144. doi:10.1093/beheco/arn125.

Dorelle, L. S., Cuña, R. H. D., Sganga, D. E., Vázquez Greco, L. L. and Nostro, F. L. L. (2020) Fluoxetine exposure disrupts food intake and energy storage in the cichlid fish Cichlasoma dimerus (Teleostei, Cichliformes). Chemosphere, 238(124609), pp. 1–8. doi:10.1016/j.chemosphere.2019.124609.

Dorelle, L. S., Cuña, R. H. D., Vázquez, G. R., Höcht, C., Zhimizu, A., Genovest, G. and Nostro, F. L. L. (2017) The SSRI fluoxetine exhibits mild effects on the reproductive axis in the cichlid fish Cichlasoma dimerus (Teleostei, Cichliformes). Chemosphere, 171, pp. 370–378. doi:10.1016/j.chemosphere.2016.11.141.

Dzieweczynski, T. L. and Hebert, O. L. (2012) Fluoxetine alters behavioral consistency of aggression and courtship in male Siamese fighting fish, Betta splendens, Physiology & Behavior, 107, pp. 92–97. doi:10.1016/j.physbeh.2012.06.007.

Eisenreich, B. R., Greene, S. and Szalda-Petree, A. (2017) Of fish and mirrors: Fluoxetine disrupts aggression and learning for social rewards. Physiology & Behavior, 173, pp. 258–262. doi: 10.1016/j.physbeh.2017.02.021.

Fent, K., Weston, A. A. and Caminada, D. (2006) Ecotoxicology of human pharmaceuticals. Aquatic Toxicology, 76, pp. 122–159. doi:10.1016/j.aquatox.2005.09.009.

Friard, O. and Gamba, M. (2016) BORIS: a free, versatile open-source event-logging software for video/audio coding and live observations. Methods in Ecology and Evolution, 7(11), pp. 1325–1330. doi: 10.1111/2041-210X.12584.

Gaworecki, K. M. and Klaine, S. J. (2008) Behavioral and biochemical responses of hybrid striped bass during and after fluoxetine exposure. Aquatic Toxicology, 88(4), pp. 207–213. doi:10.1016/j.aquatox.2008.04.011.

Genner, M. J., Turner, G. F. and Hawkins, S. J. (1999) Resource Control by Territorial Male Cichlid Fish in Lake Malawi. Journal of Animal Ecology, 68(3), pp. 522–529. doi: 10.1046/j.1365-2656.1999.00301.x.

Genner, M. J. and Turner, G. F. (2005) The mbuna cichlids of Lake Malawi: a model for rapid speciation and adaptive radiation. Fish and Fisheries, 6, pp. 1–34. doi:10.1111/j.1467-2679.2005.00173.x.

Genner, M. J., Young, K. A., Haesler, M. P. and Joyce, D. A. (2008) Indirect mate choice, direct mate choice and species recognition in a bower-building cichlid fish lek. Journal of Evolutionary Biology, 21(5), pp. 1387–1396. doi:10.1111/j.1420-9101.2008.01558.x.

Häder, D-P., Banaszak, A. T., Villafañe V. E., Narvarte, M. A., González R. A. and Helbling, E. W. (2020) Anthropogenic pollution of aquatic ecosystems: Emerging problems with global implications. Science of the Total Environment, 713(136586), pp. 1–10. doi:10.1016/j.scitotenv.2020.136586.

Hirschenhauser, K., Taborsky, M., Oliveira, T., Canàrio, A. V. M. and Oliveira, R. F. (2004) A test of the ‘challenge hypothesis’ in cichlid fish: simulated partner and territory intruder experiments. Animal Behaviour, 68, pp. 741–750. doi:10.1016/j.anbehav.2003.12.015

Holder, J. L., Barlow, G. W. and Francis, R. C. (1991) Differences in Aggressiveness in the Midas Cichlid Fish (Cichlasoma citrinellum) in Relation to Sex, Reproductive State and the Individual. Ethology, 88(4), pp. 297–306. doi:10.1111/j.1439-0310.1991.tb00284.x.

Holmstrup, M., Bindesbøl, A-M., Oostingh, G. J., Duschl, A., Scheil, V., Köhler, H-R., Loureiro, S., Soares, A. M. V. M., Ferreira, A. L. G., Kienle, C., Gerhardt, A., Laskowski, R., Kramarz, P. E., Bayley, M., Scendsen, C. and Spurgeon, D. J. (2010) Interactions between effects of environmental chemicals and natural stressors: A review. Science of the Total Environment, 408(18), pp. 3746–3762. doi:10.1016/j.scitotenv.2009.10.067.

Holzberg, S. (1978) A field and laboratory study of the behaviour and ecology of Pseudotropheus zebra (Boulenger), an endemic cichlid of Lake Malawi (Pisces; Cichlidae)M1. Journal of Zoological Systematics and Evolutionary Research, 16, pp. 171–187. doi:10.1111/j.1439-0469.1978.tb00929.x.

Hu, C.K. and Hoekstra, H.E. (2017) Peromyscus burrowing: a model system for behavioral evolution. Seminars in Cell & Developmental Biology, 61, pp. 107–114. doi:10.1016/j.semcdb.2016.08.001.

Isvaran, K. and Ponkshe, A. (2013) How general is a female mating preference for clustered males in lekking species? A meta-analysis. Animal Behaviour, 86, pp. 417–425. doi:10.1016/j.anbehav.2013.05.036.

Jaroensutasinee, M. and Jaroensutasinee, K. (2003) Type of intruder and reproductive phase influence male territorial defence in wild-caught Siamese fighting fish. Behavioural Processes, 64(1), pp. 23–29. doi:10.1016/S0376-6357(03)00106-2.

Josi, D. and Frommen, J. G. (2021) Through a glass darkly? Divergent reactions of eight Lake Tanganyika cichlid species towards their mirror image in their natural environment. Ethology, 127, pp. 925–933. doi: 10.1111/eth.13207.

Kania, B. F., Gralak, M. A. and Wielgosz, M. (2012) Four-Week Fluoxetine (SSRI) Exposure Diminishes Aggressive Behaviour of Male Siamese Fighting Fish (Betta splendens). Journal of Behavioral and Brain Science, 2, pp. 185–190. doi:10.4236/jbbs.2012.22022.

Kaufmann, P. R. and Hughes, R. M. (2006) Geomorphic and Anthropogenic Influences on Fish and Amphibians in Pacific Northwest Coastal Streams. American Fisheries Society Symposium, 48, pp. 429–455.

Kellner, M., Porseryd, T., Porsch-Hällström, I., Borg, B., Roufidou, C. and Olsén, K. H. (2018) Developmental exposure to the SSRI citalopram causes long-lasting behavioural effects in the three-spined stickleback (Gasterosteus aculeatus). Ecotoxicology, 27, pp. 12–22. doi:10.1007/s10646-017-1866-4.

Khan, I., Hou, F. and Le, H. P. (2021) The impact of natural resources, energy consumption, and population growth on environmental quality: Fresh evidence from the United States of America. Science of the Total Environment, 754(142222), pp. 1–13. doi:10.1016/j.scitotenv.2020.142222.

Kocher, T. D. (2004) Adaptive evolution and explosive speciation: the cichlid fish model. Nature, 5, pp. 288–298. doi:10.1038/nrg1316.

Kohlert, J.G., Mangan, B. P., Kodra, C., Drako, L., Long, E. and Simpson, H. (2012) Decreased Aggressive and Locomotor Behaviors in Betta Splendens after Exposure to Fluoxetine. Psychological Reports, 110(1), pp. 51–62. doi:10.2466/02.13.PR0.110.1.51-62.

Konings, A. and Kazembe, J. (2018) Nyassachromis microcephalus (errata version published in 2019). The IUCN Red List of Threatened Species 2018: e.T60990A148667319. doi:10.2305/IUCN.UK.2018-2.RLTS.T60990A148667319.en. (Accessed: 22 October 2021).

Kua, Z. X., Hamilton, I. M., McLaughlin, A. L., Brodnik, R. M., Keitzer, S. C., Gilliland, J., Hoskins, E. A., Ludsin, S. A. (2020) Water warming increases aggression in a tropical fish. Scientific Reports, 10(20107), pp. 1–13. doi:10.1038/s41598-020-76780-1.

Latifi, T., Forsatkar, M. N. and Nematollahi, M. A. (2015) Reproduction and Behavioral Responses of Convict Cichlid, Amatitlania nigrofasciata to Fluoxetine, Journal of Fisheries and Aquatic Science, 10(2), pp. 111–120. doi:10.3923/jfas.2015.111.120.

[dataset] Lee, P. (2022): Data files. figshare. Dataset. doi:10.6084/m9.figshare.21231797

Lepage, O., Larson, E. T., Mayer, I. and Winberg, S. (2005) Serotonin, but not melatonin, plays a role in shaping dominant–subordinate relationships and aggression in rainbow trout. Hormones and Behavior, 48, pp. 233–242. doi:10.1016/j.yhbeh.2005.02.012.

Lynn, S. E., Egar, J. M., Walker, B. G., Sperry, T. S. and Ramenofsky, M. (2007) Fish on Prozac: a simple, noninvasive physiology laboratory investigating the mechanisms of aggressive behavior in Betta splendens. Advances in Physiology Education, 31, pp. 358–363. doi:10.1152/advan.00024.2007.

Magalhaes, I. S., Smith, A. M. and Joyce, D. A. (2017) Quantifying mating success of territorial males and sneakers in a bower-building cichlid fish. Scientific Reports, 7(41128), pp. 1–8. doi:10.1038/srep41128.

Martin, C. H. and Genner, M. J. (2009) A Role for Male Bower Size as an Intrasexual Signal in a Lake Malawi Cichlid Fish. Behaviour, 146(7), pp. 963–978. doi:10.1163/156853908X396836.

Martin, J. M., Saaristo, M., Bertram, M. G., Lewis, P. J., Coggan, T. L., Clarke, B. O. and Wong, B. B. M. (2017) The psychoactive pollutant fluoxetine compromises antipredator behaviour in fish. Environmental Pollution, 222, pp. 592–599. doi:10.1016/j.envpol.2016.10.010.

Martínez-Ruiz, C. and Knell, R. J. (2017) Sexual selection can both increase and decrease extinction probability: reconciling demographic and evolutionary factors. Journal of Animal Ecology, 86(1), pp. 117–127. doi:10.1111/1365-2656.12601.

McKaye, k. R. (1983) Ecology and breeding behavior of a cichlid fish, Cyrtocara eucinostomus, on a large lek in Lake Malawi, Africa. Environmental Biology of Fishes, 8(2), pp. 81–96. doi:10.1007/BF00005175.

Meijide, F. J., Da Cuña, R. H., Prieto, J. P., Dorelle, L. S., Babay, P. A., and Nostro, F. L. L. (2018) Effects of waterborne exposure to the antidepressant fluoxetine on swimming, shoaling and anxiety behaviours of the mosquitofish Gambusia holbrooki. Ecotoxicology and Environmental Safety, 163, pp. 646–655. doi:10.1016/j.ecoenv.2018.07.085.

Metcalfe, C. D., Chu, S., Judt, C., Li, H., Oakes, K. D., Servos, M. Q. and Andrews, D. M. (2010) Antidepressants and their metabolites in municipal wastewater, and downstream exposure in an urban watershed. Environmental Toxicology and Chemistry, 29(1), pp. 79–89. doi:10.1002/etc.27.

Misra, A. K., Chandra, P. and Raghavendra, V. (2011) Modeling the depletion of dissolved oxygen in a lake due to algal bloom: Effect of time delay. Advances in Water Resources, 34, pp. 1232–1238. doi:10.1016/j.advwatres.2011.05.010.

Monteiro, S. C. and Boxall, A. B. A. (2010) Occurrence and Fate of Human Pharmaceuticals in the Environment. Reviews of Environmental Contamination and Toxicology, 202, pp. 53–154. doi:10.1007/978-1-4419-1157-5_2.

Moser, F. N., van Rijssel, J. C., Mwaiko, S., Meier, J. I., Ngatunga, B. and Seehausen, O. (2018) The onset of ecological diversification 50 years after colonization of a crater lake by haplochromine cichlid fishes. Proceedings of the Royal Society B, 285(20180171), pp. 1–10. doi:10.1098/rspb.2018.0171

Muschick, M., Indermaur, A. and Salzburger, W. (2012) Convergent Evolutionn within an Adaptive Radiation of Cichlid Fishes. Current Biology, 22, pp. 2362–2368. doi:10.1016/j.cub.2012.10.048.

Nielson, S. V., Kellner, M., Henriksen, P. G., Olsén, H., Hansen, S. H. and Baatrup, E. (2018) The psychoactive drug Escitalopram affects swimming behaviour and increases boldness in zebrafish (Danio rerio). Ecotoxicology, 27, pp. 485–497. doi:10.1007/s10646-018-1920-x.

Noyes, P. D., McElwee, M. K., Miller, H. D., Clark, B. W., Van Tiem, L. A., Walcott, K. C., Erwin, K. N. and Levin, E. D. (2009) The toxicology of climate change: Environmental contaminants in a warming world. Environment International, 25(6), pp. 971–986. doi:10.1016/j.envint.2009.02.006.

Parrett, J. M., Mann, D. J., Chung, A. Y. C., Slade, E. M. and Knell, R. J. (2019) Sexual selection predicts the persistence of populations within altered environments. Ecology Letters, 22(10), pp. 1629–1637. doi:10.1111/ele.13358.

Paterson, G., and Metcalfe, C. D. (2008). Uptake and depuration of the anti-depressant fluoxetine by the Japanese medaka (Oryzias latipes). Chemosphere, 74(1), pp. 125–130. doi:10.1016/j.chemosphere.2008.08.022.

Pauers, M. J., Kapfer, J. M., Fendos, C. E. and Berg, C. S. (2008) Aggressive biases towards similarly coloured males in Lake Malawi cichlid fishes. Biology Letters, 4(2), pp. 156–159. doi:10.1098/rsbl.2007.0581.

Perreault, H. A. N., Semsar, K. and Godwin, J. (2003) Fluoxetine treatment decreases territorial aggression in a coral reef fish, Physiology & Behavior, 79, pp. 719–724. doi:10.1016/S0031-9384(03)00211-7.

Polverino, G., Martin, J. M., Bertram, M. G., Soman, V. R., Tan, H., Brand, J. A., Mason, R. T. and Wong, B. B. M. (2021) Psychoactive pollution suppresses individual differences in fish behaviour. Proceedings of the Royal Society B, 288(20202294), pp. 1–9. doi:10.1098/rspb.2020.2294.

Prasad, P., Ogawa, S. and Parhar, I. S. (2015) Role of serotonin in fish reproduction, Frontiers in Neuroscience, 9(195), pp. 1–9. doi:10.3389/fnins.2015.00195.

Random.org (2021a) True Random Number Service. Available at: https://www.random.org/ (Accessed: 7 October 2021).

Random.org (2021b) Random Sequence Generator. Available at: https://www.random.org/sequences/ (Accessed: 7 October 2021).

Ratnasabapathi, Dr. D. and Souchek, J. B. R. (1992) Effects of temperature and prior residence on territorial aggression in the convict cichlid Cichlasoma nigrofasciatum. Aggressive Behavior, 18(5), pp. 365–372. doi:10.1002/1098-2337(1992)18:5<365::AID-AB2480180506>3.0.CO;2-E.

Roggatz, C. C., Fletcher, N., Benoit, D. M., Algar, A. C., Doroff, A., Wright, B., Wollenberg Valero, K. C. and Hardege, J. D. (2019) Saxitoxin and tetrodotoxin bioavailability increases in future oceans. Nature Climate Change, 9, pp.840–844. doi:10.1038/s41558-019-0589-3.

Rowe, L. and Houle, D. (1996) The Lek Paradox and the Capture of Genetic Variance by Condition Dependent Traits. Proceedings of the Royal Society B, 263(1375), pp. 1415–1421. doi:10.1098/rspb.1996.0207.

RStudio Team (2020) RStudio: Integrated Development Environment for R. RStudio, PBC, Boston, MA. Available at:m! http://www.rstudio.com/ (Accessed: 7 October 2021).

Ruhoy, I. S. and Daughton, C. G. (2008) Beyond the medicine cabinet: An analysis of where and why medications accumulate, Environment International, 34, pp. 1157–1169. doi:10.1016/j.envint.2008.05.002.

Saaristo, M., Brodin, T., Balshine, S., Bertram, M. G., Brooks, B. W., Ehlman, S. M., McCallum, E. S., Sih, A., Sundin, J., Wong, B. B. M. and Arnold, K. E. (2018) Direct and indirect effects of chemical contaminants on the behaviour, ecology and evolution of wildlife. Proceedings of the Royal Society B, 285(20181297), pp. 1–10. doi:10.1098/rspb.2018.1297.

Secretariat of the Convention on Biological Diversity (2014) Global Biodiversity Outlook 4. Montréal, Canada. Available at: https://www.cbd.int/gbo/gbo4/publication/gbo4-en.pdf (Accessed: 18 October 2021).

Schaedelin, F. C. and Taborsky, M. (2006) Mating craters of Cyathopharynx furcifer (Cichlidae) are individually specific, extended phenotypes. Animal Behaviour, 72(4), pp. 753–761. doi:10.1016/j.anbehav.2005.11.028.

Seehausen, O. (2000) Explosive Speciation Rates and Unusual Species Richness in Haplochromine Cichlid Fishes: Effects of Sexual Selection, Advances in Ecological Research, 31, pp.237–274. doi:10.1016/S0065-2504(00)31015-7.

Seehausen, O. (2006) African cichlid fish: a model system in adaptive radiation research, Proceedings of the Royal Society B, 273, pp. 1987–1998. doi:10.1098/rspb.2006.3539.

Seehausen, O. and van Alphen, J. J. M. (1999) Can sympatric speciation by disruptive sexual selection explain rapid evolution of cichlid diversity in Lake Victoria? Ecology Letters, 2, pp. 262–271.

Seehausen, O. and Schluter, D. (2004) Male–male competition and nuptial-colour displacement as a diversifying force in Lake Victoria cichlid fishes. Proceedings of the Royal Society of London B, 271, pp. 1345–1353. doi:10.1098/rspb.2004.2737.

Selz, O. M., Pierotti, M. E. R., Maan, M. E., Schmid, C. and Seehausena, O. (2014) Female preference for male color is necessary and sufficient for assortative mating in 2 cichlid sister species. Behavioral Ecology, 25(3), pp. 612–626. doi:10.1093/beheco/aru024.

Selz, O. M., Thommen, R., Pierotti, M. E. R., Anaya-Rojas, J. M. and Seehausen, O. (2016) Differences in male coloration are predicted by divergent sexual selection between populations of a cichlid fish. Proceedings of the Royal Society B, 283(20160172), pp. 1–9. doi:10.1098/rspb.2016.0172.

Schwindt, A. R., Winkelman, D. L., Keteles, K., Murphy, M. and Vajda, A. M. (2014) An environmental oestrogen disrupts fish population dynamics through direct and transgenerational effects on survival and fecundity. Journal of Applied Ecology, 51, pp. 582–591. doi:10.1111/1365-2664.12237.

Shih, S. (2017) Effects of Fluoxetine on Social and Startle Behavior in the African Cichlid Astatotilapia burtoni, MA thesis, CUNY Hunter College, New York. Available at: https://academicworks.cuny.edu/cgi/viewcontent.cgi?article=1268&context=hc_sas_etds. (Accessed: 22 October 2021)

Silva, L. J. G., Lino, C. M., Meisel, L. M. and Pena, A. (2012) Selective serotonin re-uptake inhibitors (SSRIs) in the aquatic environment: An ecopharmacovigilance approach. Science of the Total Environment, 437, pp. 185–195. doi:10.1016/j.scitotenv.2012.08.021.

Stahl, S. M. (1998) Mechanism of action of serotonin selective reuptake inhibitors: Serotonin receptors and pathways mediate therapeutic effects and side effects. Journal of Affective Disorders, 51(3), pp. 215–235. doi:10.1016/S0165-0327(98)00221-3.

Stauffer Jr., J. R. and Gray, E. V. S. (2004) Phenotypic plasticity: its role in trophic radiation and explosive speciation in cichlids (Teleostei: Cichlidae). Animal Biology, 54(2), pp. 137–158. doi:10.1163/1570756041445191.

Stauffer Jr., J. R., McKaye, K. R. and Konings, A. F. (2002) Behaviour: an important diagnostic tool for Lake Malawi cichlids. Fish and Fisheries, 3, pp. 213–224.

Srivastava, K. (2009) Urbanization and mental health, Industrial Psychiatry Journal, 18(2), pp. 75–76. doi: 10.4103/0972-6748.64028.

Sumpter, J. P., Donnachie, R. L. and Johnson, A. C. (2014) The apparently very variable potency of the anti-depressant fluoxetine. Aquatic Toxicology, 151, pp. 57–60. doi:10.1016/j.aquatox.2013.12.010.

Taylor, M. I., Turner, G. F., Robinson, R. L. and Stauffer Jr, J. R. (1998) Sexual selection, parasites and bower height skew in a bower-building cichlid fish, Animal Behaviour, 56(2), pp. 379–384. doi:10.1006/anbe.1998.0795.

Trewavas, E. (1935) A synopsis of the cichlid fishes of Lake Nyasa. Annals and Magazine of Natural History Series 10, 16(91), pp. 65–118. doi:10.1080/00222933508655026

Turner, G. F., Seehausen, O., Knight, M. E., Allender, C. J. and Robinson, R. L. (2001) How many species of cichlid fishes are there in African lakes? Molecular Ecology, 10, pp. 793–806.

Walker II, W. H., Meléndez-Fernández, O. H., Nelson, R. J. and Reiter, R. J. (2019) Global climate change and invariable photoperiods: A mismatch that jeopardizes animal fitness. Ecology and Evolution, 9, pp. 10044–10054. doi:10.1002/ece3.5537.

Wilkinson, J. L., Boxall, A. B. A., Kolpin, D. W., Leung, K. M. Y., Lai, R. W. S., Galbán-Malagón, C., Adell, A. D., Mondon, J., Metian, M., Marchant, R. A., Bouzas-Monroy, A., Cuni-Sanchez, A., Coors, A., Carriquiriborde, P., Rojo, M., Gordon, C., Cara, M., Moermond, M., Luarte, T., Petrosyan, V., Perikhanyan, Y., Mahon, C. S., Mcgurk, C. J., Hofmann, T., Kormoker, T., Iniguez, V., Guzman-Otazo, J., Tavares, J. L., Gildasio De Figueiredo, F., Razzolini, M. T. P., Dougnon, V., Gbaguidi, G., Traoré, O., Blais, J. M., Kimpe, L. E., Wong, M., Wong, D., Ntchantcho, R., Pizarro, J., Ying, G., Chen, C., Páez, M., Martínez-Lara, J., Otamonga, J., Poté, J., Ifo, S. A., Wilson, P., Echeverría-Sáenz, S., Udikovic-Kolic, N., Milakovic, M., Fatta-Kassinos, D., Ioannou-Ttofa, L., Belušová, V., Vymazal, J., Cárdenas-Bustamante, M., Kassa, B. A., Garric, J., Chaumot, A., Gibba, P., Kunchulia, I., Seidensticker, S., Lyberatos, G., Halldórsson, H. P., Melling, M., Shashidhar, T., Lamba, M., Nastiti, A., Supriatin, A., Pourang, N., Abedini, A., Abdullah, O., Gharbia, S. S., Pilla, F., Chefetz, B., Topaz, T., Yao, K. M., Aubakirova, B., Beisenova, R., Olaka, L., Mulu, J. K., Chatanga, P., Ntuli, V., Blama, N. T., Sherif, S., Aris, A. Z., Looi, L. J., Niang, M., Traore, S. T., Oldenkamp, R., Ogunbanwo, O., Ashfaq, M., Iqbal, M., Abdeen, Z., O’dea, A., Morales-Saldaña, J. M., Custodio, M., De La Cruz, H., Navarrete, I., Carvalho, F., Gogra, A. B., Koroma, B. M., Cerkvenik-Flajs, V., Gombač, M., Thwala, M., Choi, K., Kang, H., Ladu, J. L. C., Rico, A., Amerasinghe, P., Sobek, A., Horlitz, G., Zenker, A. K., King, A. C., Jiang, J., Kariuki, R., Tumbo, M., Tezel, U., Onay, T. T., Lejju, J. B., Vystavna, Y., Vergeles, Y., Heinzen, H., Pérez-Parada, A., Sims, D. B., Figy, M., Good, D. and Teta, C. (2022) Pharmaceutical pollution of the world’s rivers. PNAS, 119(8), pp. 1–10. doi:10.1073/pnas.211394711.

Winder, V. L., Sapozhnikova, Y., Pennington, P. L., and Wirth, E. F. (2009) Effects of fluoxetine exposure on serotonin-related activity in the sheepshead minnow (Cyprinodon variegatus) using LC/MS/MS detection and quantitation. Comparative Biochemistry and Physiology, Part C, 149(4), pp. 559–565. doi:10.1016/j.cbpc.2008.12.008.

Young, K. A., Genner, M. J., Joyce, D. A. and Haesler, M. P. (2009) Hotshots, hot spots, and female preference: exploring lek formation models with a bower-building cichlid fish. Behavioral Ecology, 20(3), pp. 609–615. doi:10.1093/beheco/arp038.

Zubizarreta, L., Perrone, R., Stoddard, P. K., Costa, G. and Silva, A. C. (2012) Differential serotonergic modulation of two types of aggression in weakly electric fish. Frontiers in Behavioral Neuroscience, 6(77), pp. 1–10. doi:10.3389/fnbeh.2012.00077.

